# Integrated management of Canada thistle (*Cirsium arvense*) in the Great Plains and Intermountain West using a biocontrol agent (*Puccinia suaveolens)*

**DOI:** 10.1101/2025.03.19.644225

**Authors:** Caitlin Henderson, Kristi Gladem, Stephen L. Young, Dan W. Bean, Robert N. Schaeffer

**Author notes:** Authors for correspondence: **Caitlin Henderson; Email:****, Kristi Gladem; Email:**. These authors contributed equally to the study (Caitlin Henderson, Kristi Gladem).

## Abstract

Canada thistle [*Cirsium arvense* (L.) Scop.] is an invasive perennial plant that threatens agricultural landscapes and natural ecosystems worldwide. The extensive regenerative root system of Canada thistle complicates control efforts, with current strategies having limited success. *Puccinia suaveolens* (Pers.) Rostr (syn. *Puccinia punctiformis* (F. Strauss) Rohl), an obligate biotrophic rust fungus, has shown potential as a biological control agent by systemically infecting the root system, reducing root mass and shoot growth, and limiting vegetative regeneration; however, its efficacy when integrated with other control methods remains unclear. We conducted experiments from 2020 to 2022 at two sites in Colorado and Utah to evaluate *P. suaveolens* efficacy when applied alone and in combination with mowing, tillage, and herbicide. Treatments were applied in Fall (2020 and 2021), with monitoring of thistle stem density, vegetative cover, as well as *P. suaveolens* incidence before and after treatments through 2022. While *P. suaveolens* alone contributed to a decrease in thistle density, it was less effective compared to herbicide treatments, and its impact when integrated with mowing or tillage was inconsistent. Herbicide application (alone and when combined with *P. suaveolens*) generated the greatest immediate reduction in thistle stem density and vegetative cover, although it resulted in the greatest amount of bare ground exposure. Grass coverage present within treated plots varied significantly between treatments, ranging from 0-75%, with the highest percentage observed in herbicide treatments compared to control and tillage in both years. Forb cover remained below 10% across treatments and years. Although *P. suaveolens* continues to show promise as a biological control agent, further research is needed to improve efficacy, and optimize integration with other control strategies.

## Introduction

*Cirsium arvense* (Asteraceae), commonly referred to as Canada thistle, is a pervasive perennial weed found throughout temperate regions globally. In the Western United States (U.S.), Canada thistle ranks as one of the highest among the most commonly occurring noxious weeds, posing significant threats to both managed and natural landscapes (Bodo Slotta et al. 2010; Moore et al. 1975; Nuzzo 1997). In agricultural systems, Canada thistle competes for light, nutrients, and water, leading to reduced crop yield and quality, generating significant economic losses for producers (Jacobs et al. 2006; Moyer et al. 1991; O’Sullivan 1982). In natural systems, it similarly competes with and displaces native plant species (Jacobs 2006). Canada thistle is commonly found in disturbed areas, including roadsides, streambanks, ditches, clear cuts, forest openings, and wet or wet-mesic grasslands and rangelands, as part of the initial post-disturbance community (Morishita 1999; Nuzzi 1997). It is prevalent in nearly every upland herbaceous community within its range, particularly prairie communities and riparian habitats (Nuzzo 1997).

Canada thistle survives and spreads through two reproductive strategies: sexual reproduction via seeds and clonal vegetative growth. Seeds are small and light, with highly variable germination success (Hodgson 1964; Moore 1975; Nuzzo 1997); however seeds are important for range expansion as shown by the genetic diversity of North American populations (Bodo Slotta et al 2010). Once established, the plant develops a creeping root system, up to 2-3m belowground, with adventitious root buds resulting in clonal vegetative growth that enables rapid propagation and spread (Donald 1994; Lalonde and Roitberg 1994). The adventitious buds develop into new rosettes and lateral roots that continue to grow throughout spring, summer, and into fall. As temperatures decrease in fall, the aboveground vegetation dies off, and the roots overwinter; in the spring, root growth resumes and new shoots emerge (Lalonde and Roitberg 1994; Tiley 2010). Fragments of roots as small as 1cm are capable of regenerating and forming new colonies (Nadeau and Vanden Born 1989; Thomsen et al. 2015). The complex and extensive root system of Canada thistle allows for propagation, spread, and recovery making it particularly problematic and challenging for management (Nadeau and Vanden Born 1989; Tiley 2010).

A number of management tactics are regularly utilized in efforts to control Canada thistle. Chemical control is common and effective, generally providing rapid results. Mowing can also be an effective control method by reducing photosynthetic capacity aboveground and depleting root reserves used for regrowth, leading to a reduction of new shoots the following season (Bourdôt et al. 2011; Graglia et al. 2006). Similarly, cultivation and tillage fragment the root system and force the plant to use root reserves for recovery, however this may also promote new shoot development and further spread of Canada thistle (Graglia et al. 2006, Thomsen et al. 2015). Both tillage and mowing are most effective when integrated into a weed management program to control established populations. While most current management of Canada thistle relies on either herbicides, mowing, or cultivation, these tactics can be costly, labor-intensive, and often not ideal for environmentally sensitive areas (e.g., riparian zones) (Bourdôt et al. 2011; Graglia et al. 2006; Peterson et al. 2020; Thomsen et al. 2015). In contrast, biological control tactics are often more suitable for managing weeds in natural areas, as they can be self-perpetuating, and more economical in landscape-wide suppression of target species (Cripps et al. 2011; Guske et al. 2004; Peterson et al. 2020; Sciegienka et al. 2011).

Effective biological control has long been sought for Canada thistle. *Puccinia suaveolens* (Pers.) Rostr (syn. *Puccinia punctiformis* (F. Strauss) Rohl), an obligate biotrophic rust fungus, was first proposed as a control method for Canada thistle in 1893 in North America (Wilson 1969). *Puccinia suaveolens* can be naturally found on Canada thistle plants throughout its growing region, commonly co-introduced in the invasive range and causing periodic outbreaks of disease (Berner et al. 2013; French and Lightfield 1990). Highly host specific to Canada thistle, *P. suaveolens* has only been reported on two other thistles native to Eurasia; *Silybum marianum* (L.) Gaertner in 2002 under greenhouse conditions and *C. setosum* (Willd.) M. Bieb. in 2023 under field conditions in China (Berner et al. 2002; Liang et al. 2024). The potential for *P. suaveolens* to be utilized more broadly as a biocontrol for Canada thistle will increase with continued research (Bean et al. 2024; Berner et al. 2015a; Berner et al. 2015b; Cripps et al. 2014; Thomas et al. 1994).

The lifecycle of *P. suaveolens* can be divided in two stages: the vegetative mycelium within the root system and the spore-producing aboveground systemic and local infections. It is thought that *P. suaveolens* remains latent within the root system until abiotic or biotic conditions are adequate to produce spore-bearing systemically infected stems (Mendgen and Hahn, 2002). *Puccinia suaveolens* infection reduces Canada thistle belowground biomass as root resources are parasitized by mycelia and also allocated to plant defense compounds instead of growth (Chichinsky et al., 2023; Clark et al. 2020). This reduces the aboveground shoots and competitive ability of Canada thistle (Chichinsky et al., 2023). Systemically infected stems typically do not flower, can die off early in the season and may help to further reduce root resources of infected Canada thistle colonies (Van den Ende et al. 1987; Chichinsky et al. 2023). The rate and distance of spread of *P. suaveolens* caused by underground mycelia or aboveground spores remains unknown.

*Puccinia suaveolens* produces five spore types that develop consecutively beginning with emergence of systemically infected shoots in early spring that are deformed, chlorotic, strongly floral scented, and covered in yellow-orange pycnia pustules (Buller 1950; Menzies 1953; Petersen 1974). After outcrossing, pycniospores give rise to chain-like formations of the dark orange-brown aeciospores giving the systemically infected stems a characteristic rusty appearance (Berner et al. 2013; Connick and French et al. 1991). Production of urediniospores is indicated by the darkening of the spores (Buller 1950; Peterson 1974). Urediniospores and aeciospores are morphologically indistinguishable as single celled spores but only urediniospores produce localized infections on neighboring plants throughout summer (Kirk et al. 2001, Menzies 1953, Peterson 1974). Localized infections produce pustules on the leaves but the shoots do not have the same growth abnormalities associated with systemic infection, and will still appear relatively normal (Baka and Lösel 1992; Thomas et al. 1994). Localized infections can produce two-celled teliospores on leaves that senesce heading into fall, then the spores will either blow off and overwinter in the soil or germinate on new rosettes to initiate new vegetative infection in the roots or systemic infection aboveground. (Alexopoulos et al. 1996; Berner et al. 2015b; Menzies 1953). Optimal teliospore germination occurs when temperatures are between 16-20°C (Berner et al. 2013; French and Lightfield 1990) with optimal dew periods of 2-3 hours (Morin et al. 1992a, 1992b).

Integrated weed management (IWM) is a holistic approach implementing one or more biological, physical, cultural or chemical control tactics (Harker and Donovan 2013). IWM aims to reduce weed adaptation and resistance to any single control tactic by using several possible tactics that take into account threshold populations, critical periods, and environmental outcomes. Utilizing IWM may lead to reduced environmental impacts of any given control method, decreased control costs by reducing pests to economically and ecologically insignificant levels, increased sustainability and reduced herbicide resistance (Harker and Donovon 2013). In a meta-analysis, Davis et al. (2018) found that combined treatments had better long-term outcomes for control of Canada thistle then reliance on herbicide treatment alone. Mowing and tillage have been shown to affect Canada thistle populations, but results have varied between significant reductions in population size to virtually no impact at all (Beck and Sebastian 2000; Bourdôt et al. 2011). A stem-mining weevil, *Hadroplontus litura,* and a bacterial plant pathogen, *Pseudomonas syringae* pv. *tagetis,* have shown to have an additive effect in suppressing Canada thistle treated with herbicide (Sciegienka 2011). *Puccinia suaveolens* has also previously been used in conjunction with other control methods. Mowing combined with *P. suaveolens* strongly reduced the proportion of fertile flower heads of Canada thistle compared to infection alone (Kluth et al. 2003). Demers et al. (2006) found that systemically infected *P. suaveolens* shoots increased, while healthy shoots decreased when combined with mowing. In a greenhouse experiment with a crop sequence of winter wheat, spring pea and summer safflower, crop competition reduced Canada thistle but when inoculated with *P. suaveolens*, the effect was increased (Chichinsky et al 2023). The potential of using *P. suaveolens* in an IWM approach for Canada thistle is supported, but more research is needed to develop and refine best management practices.

Canada thistle is a problematic weed that is difficult to control and current methods have varying degrees of success in both managed and natural ecosystems. As a biological control for Canada thistle, *P. suaveolens* has significant potential, as it can self-perpetuate, spread to surrounding areas, and contribute to population suppression at large scales when applied alone (Bean et al. 2024). Our objectives were to determine the efficacy and compatibility of different control methods (mowing, tillage, herbicide, and *P. suaveolens*) when applied alone and in combination to suppress Canada thistle. We highlight the benefits and limitations of using *P. suaveolens* in an IWM program, along with considerations for improved application efficacy.

## Materials and Methods

### Study Sites

Two experimental sites were established in 2020: one in the Tamarack Ranch State Wildlife Area of Colorado (CO, 40.8320°N, 102.80437°W) and the other in Park City, Utah (UT 40.674330°N, 111.491324°W). The CO site is within the High Plains Ecoregion, while the UT site is within the Wasatch and Uinta Range Ecoregion (Omernik and Griffith 2014). The CO site is a 12 by 600 m plot of land, characterized as a shortgrass prairie with seasonal water inundation. Historically, the site had been maintained as a food crop plot for wildlife. At the end of each growing season, glyphosate had been applied and the plot was tilled for several years, resulting in the formation of a near monoculture of Canada thistle (Levi Kokes, personal communication May 2020). The CO site typically has precipitation occurring throughout the year (Supplementary Table S:1). The UT site is a small preserve nestled within a suburban development. Historically, spraying, particularly for musk thistle (*Carduus nutans* L.), and goat/sheep grazing had been occasionally employed at the UT site but had not occurred for many years; the area is largely left untouched (Sara Jo Dickens, personal communication, 2020). The UT site generally has the majority of precipitation occurring in the winter months (S:1) (PRISM Climate Group, 2023).

### Puccinia suaveolens Inoculum

Dried inoculum was prepared following Berner et al (2013) and Bean et al (2024). Briefly, Canada thistle leaves bearing telia (small pustules on yellowing leaves), were collected in late summer from a site near Colorado Springs, CO. Leaves were harvested and stored in paper bags to allow foliage to dry at room temperature. Dried leaves were ground to a coarse powder in a kitchen blender and used as inoculum within season, or stored at −80 ◦C for future use. Samples of ground leaf preparations were viewed under a microscope to ensure the majority of spores were two-celled teliospores, which are necessary for initiating systemic infection (French and Lightfield, 1990; Berner et al 2013; Van Den Ende et al. 1987).

### Experimental Design

In both UT and CO, an experiment site was established using a randomized complete block design, consisting of 10 treatment combinations applied across replicates. Treatments included an untreated control, *P. suaveolens* inoculation alone, tillage, tillage plus *P. suaveolens* inoculation, mowing, mowing plus *P. suaveolens* inoculation, herbicide (aminopyralid and chlorsulfuron tank mix), herbicide plus *P. suaveolens* inoculation, herbicide, mowing, and tillage (HMT), and HMT plus *P. suaveolens* inoculation (Table 1). Each treatment was applied once in 2020 and 2021 to field plots (CO: 2 by 6 m; UT: 2 by 5 m). Plots were spaced (CO:4m; UT:2m) apart to avoid edge effects with 8 replicates in CO and 4 in UT. Differences in experimental set up between sites were due to the size and accessibility of Canada thistle populations.

Herbicide and mowing treatments were applied in the fall during the first week of September. An herbicide tank mix was applied (aminopyralid 7 fl. oz/acre; chlorsulfuron at 1 fl. oz/acre) using a backpack sprayer calibrated in the field. Aminopyralid was chosen specifically as it is more effective at lower rates compared to other herbicides (e.g., picloram and clopyralid), and may also be used in areas where other chemicals are not appropriate or recommended (Enloe et al. 2017). In herbicide and mowing combination treatments, mowing was applied first to provide an opportunity for more even herbicide application and uptake given the physical damage to Canada thistle (Carpinelli 2004). Fourteen days after initial treatments with mowing and herbicide, tillage (30 cm depth) and *P. suaveolens* inoculum (40 g CO, 33.3 g UT) were applied to select plots. When applying *P. suaveolens* inoculum, the entire plot was first sprayed with water using a backpack sprayer to create a mist on Canada thistle leaves, then *P. suaveolens* was broadcast by hand no higher than 1m above the ground to avoid excessive dispersal by wind. The 14-day period allowed the herbicide to translocate through the roots and other tissues before tillage following recommended manufacturer (Corteva) guidelines. *Puccinia suaveolens* inoculum was applied last either alone or in combination (Table 1). The later timing for inoculum application in IWM treatments allowed for new growth and rosettes of Canada thistle in response to mowing and possibly tillage, which may improve the chance for infection (Demers et al. 2006). Applications of *P. suaveolens* inoculum before mowing, tillage, or herbicide spray, would have been detrimental to germinating teliospores, which may have begun developing mycelia in the live tissue and subsequently been destroyed (Berner et al. 2013; Petersen 1974).

The initial monitoring of plots at both sites occurred in fall prior to first treatments. Monitoring occurred two weeks prior to the optimal timing for *P. suaveolens* teliospore inoculum application at each respective site. At both sites, a quadrat (m^2^) was placed in the center of each plot and the number of thistle stems was counted and percent groundcover of Canada thistle, grass, forbs, litter, and bare ground was estimated. Across the entire plot, a two-minute timed count of Canada thistle stems systemically infected with *P. suaveolens* was also performed.

### Statistical Analyses

All analyses were performed with R (R Core Team 2023), using packages tidyverse, ggplot2, glmmTMB, DHARMa, emmeans and car (Brooks et al. 2017; Fox and Weisberg 2019; Hartig 2022; Lenth 2024; Wickham et al. 2019). Data from the two sites were analyzed separately because of the imbalance in design due to the difference in site accessibility and size of Canada thistle population. Stem density change as a function of *P. suaveolens* inoculum application (Yes or No); management approach [Control, Herbicide (H), Mowing (M), Tillage (T), and HMT]; or year (Fall 2020, Fall 2021, and Fall 2022); and the interaction between combinations of these parameters were analyzed with a generalized linear model (GLM). Stem density was modeled (with negative binomial) also using GLM. Significance was tested using ANOVA type II Wald chi-square tests, followed by post-hoc pairwise Tukey test. The percent change in stem density was calculated with standard deviation. Finally, cover data were analyzed using a GLM (with beta distribution) testing significance with ANOVA type II Wald chi-square tests, followed by post-hoc pairwise Tukey test.

## Results and Discussion

In this study, conducted in two regions of the western U.S., the Intermountain West and the Great Plains, we evaluated the efficacy of *P. suaveolens* and its compatibility with other control methods for managing Canada thistle by measuring stem density and vegetative cover. While stem density reflects the direct effects on the target weed, vegetative cover can represent biodiversity, forage availability, resiliency of the landscape, nutrient cycling, and be used to predict production costs for livestock producers or fire risk. Consideration of both stem density and resulting vegetative cover will help land managers to make informed decisions about which treatments work in their IWM plan and how *P. suaveolens* can be incorporated.

### Stem Density

Stem density of Canada thistle differed significantly between treatments (UT: P<0.0001, CO: P<0.0001) (Figure 1, Table 3). Herbicide treatments resulted in the largest decrease in stem density, while tillage treatments had the largest increase in stem density. There was also a significant interaction between management and year (UT: P<0.0001, CO: P<0.0001), and a marginal interaction between management, season, and *P. suaveolens* in CO (P=0.057) (Table 3). At the UT site, initial average stem density was 24 ± 3 shoots per m^2^ (Figure 1), while the CO site initial average stem density was 80 ± 36 shoots per m^2^ (Figure 1). These differences may be attributed to variation in climate conditions and prior management practices employed at each site. During post-treatment monitoring, stem density in the control plots at the UT site increased by 85% (stem count m^2^ (sc), 2020: 22±16; 2021: 40±9; 2022: 43±19) the first year, with an additional 7% increase the following year (Table 2 and Figure 1). At the CO site, stem density in the control plots decreased by 3% (sc, 2020: 76±31; 2021: 74±28; 2022: 63±55) in the first year and then decreased another 15% the following year (Table 2 and Figure 1).

The UT site has higher average annual precipitation compared to the CO site, although during the first year of treatments they were about equal (Supplementary Table S1). At the UT site, most precipitation occurs as winter snowfall, resulting in extended dry periods that can stress Canada thistle, reducing root and stem growth, and subsequently, stem density and coverage (Tiley 2010). In contrast, Tamarack Ranch, CO, receives more evenly distributed precipitation throughout the year (PRISM Climate Group 2023, Supplementary Table S1). At the UT site, temperature ranged from −11.1°C during the coldest months to 29°C in the hottest months of the experiment period (2020-2022). At the CO site, the temperature ranged from −13.5°C during the coldest months to 33°C (2020-2022) during the hottest months (Supplementary Table S1) (PRISM Climate Group 2023).

### Herbicide and Herbicide+Mowing+Tillage (HMT)

Herbicide treatments, whether alone or in combination, were most effective in decreasing Canada thistle stem density, and were significantly different compared to the other treatments (UT: P<0.001; CO: P<0.001; Table 3). At both sites, there was an immediate decline in stem density that continued even after the second application with sparse regrowth (Figure 1). In CO, herbicide only treatments decreased stem density 93% in year one and another 77% in year two (sc, 2020: 78±21; 2021: 6±6; 2022: 1±2). When *P. suaveolens* was applied along with the herbicide, stem density declined 90% in year one and 100% the following year (sc, 2020: 84±40; 2021: 8±9; 2022: 0±0). The UT site had similar decreases in stem density from herbicide, year one 97% and year two 100% (sc, 2020: 29±13; 2021: 1±2; 2022: 0±0) and herbicide with *P. suaveolens*, year one 98% and year two 100% (sc, 2020: 22±14; 2021: 1±1; 2022: 0±0) (Table 2).

Using Tukey’s fence method for determining outliers, the Colorado HMT and HMT with *P. suaveolens* plots in block 8 in 2022 were determined to be outliers and were removed from analysis. Deep rooted perennial grasses that may provide higher competition were disturbed by tillage and replaced with annual grasses. Herbicide effects were perhaps lessened by seasonal water inundation. However the explanation of these outliers is unknown and further research would be needed to confirm the potential cause of increased Canada thistle stem density in these plots.

In CO, the combined treatment (HMT) without *P. suaveolens* had a 97% decrease in stem density in year one and a 93% decrease in year two (sc, 2020: 67±44; 2021: 2±4; 2022: 0±0). When *P. suaveolens* was applied along with the HMT treatment, stem density decreased 86% in year one and 94% in year two (sc, 2020: 84±47; 2021: 12±15; 2022: 1±1) (Table 2). At the UT site, stem density in the HMT plots without *P. suaveolens* decreased 69% in year one and another 95% in year two (sc, 2020: 25±10; 2021: 8±5; 2022: 1±3). When *P. suaveolens* was applied along with the HMT treatment, stem density decreased 84% in year one and 100% in year two (sc, 2020: 27±7; 2021: 4±5; 2022: 0±0) (Table 2). Aminopyralid, which is selective against broadleaf weeds in rangelands and pastures, provided near 100% control of Canada thistle in herbicide-treated plots with additive effects from *P. suaveolens* inoculum application. Limited thistle regeneration was observed, likely emanating from neighboring plants, seeds, or remaining root fragments.

### Puccinia suaveolens

*Puccinia suaveolens* was present at both sites in plots after treatments, however, there were only a few symptomatic stems. In CO, no symptomatic plants were found during the fall monitoring. In UT, the symptomatic shoots were found in the tillage plus *P. suaveolens* treatment, first appearing in year one (1 shoot) and also in year two (4 shoots). There was an overall lack of statistical significance (Table 3) of *P. suaveolens* application at both sites, however there was a general declining trend in stem density indicating that the *P. suaveolens* had a slight suppressing effect on Canada thistle (Figure 1, Table 2).

In UT, *P. suaveolens* treatments alone appeared to slow the increase of stem density (48%) by year two (sc, 2020: 26±8; 2022: 38±13) when compared to the untreated control, which had an increase of 98% (sc, 2020: 22±16; 2022: 43±19) (Table 2). In CO, *P. suaveolens* treatment (sc, 2020: 93±28; 2022: 54±35), reduced stem density 24% more than the untreated control (sc, 2020: 76±31; 2022: 63±55) (Table 2). The decrease in Canada thistle stem density following *P. suaveolens* is similar to the findings of Bean et al. (2024), who recorded stem density decreases at 77% of monitored sites in Colorado over 3-8 years that went from 87.9 ± 6.5 stems to 44.7 ± 4.2 stems. Sites with more frequent and higher quantities of *P. suaveolens* inoculum applied had a lower stem density over time. We suspect that Canada thistle stem densities within *P. suaveolens* treated plots will continue to decrease with or without additional inoculations.

While stem decline was observed, the lack of symptomatic thistle stems could potentially be attributed to genotypic differences and associated resistance within Canada thistle, or the compatibility of the host-pathogen interaction. Alternatively the abiotic or biotic factors that induce production of systemically infected stems may not have been met though vegetative mycelium within the root system could still be present. *Puccinia suaveolens* may continue to have an impact on Canada thistle or additional inoculation treatments might be required. This could be the case at both sites, but particularly at the CO site, where stem density decline was more obvious in Fall 2022 after a second inoculation (Figure 1).

### Mowing

A slight interaction occurred between season and *P. suaveolens* in CO (P=0.058), the significance occurring in the second year (P=0.0116) with mowing plus *P. suaveolens* having a greater decrease in stem density (65%) (sc, 2020: 62±41; 2022: 22±25) compared to mowing alone (49%) (sc, 2020: 83±37; 2022: 43±35). In UT, mowing (sc, 2020: 22±9; 2022: 21±11) and mowing plus *P. suaveolens* (sc, 2020: 25±17; 2022: 22±8) resulted in a small decrease in stem density of 1% and 13% respectively with no significance.

The reduction in stem density between mowing with *P. suaveolens* inoculation and mowing alone was not statistically significant in UT and only slightly in CO (P=0.099, Table 3). In CO, mowing with *P. suaveolens* inoculation initially had a smaller impact compared to mowing alone. However, in year two mowing with inoculations showed a slight decrease in stem density compared to mowing alone, which had a slight increase in stem density (Figure 1) (Table 3). In UT, Canada thistle stem density resulting from mowing (averaged over *P. suaveolens*) was significantly lower compared to control (averaged over *P. suaveolens*), (UT: P=0.003, CO P=0.005) (Table 3). In UT, mowing had significantly lower stem density (P=0.006) compared to tillage, with no significance in CO (Table 2). Mowing has been used to enhance the occurrence of systematically infected stems (Bourdôt et al. 2011), and increases localized infection by spreading spores (Demers et al. 2006). Very few systemically infected stems were found during the 2-year study, which may have resulted in less additive effects from mowing with *P. suaveolens* compared to mowing alone. However, mowing should still be utilized with *P. suaveolens* in an IWM program, as the two treatments can be compatible and mutually beneficial.

### Tillage

In CO, there was a significant difference (P=0.043, Table 3) between tillage and control. Further analysis showed that tillage treatments had significantly greater decline in stem density compared to control in 2022 (P=0.003). There was no significant difference between tillage alone and tillage with *P. suaveolens* (UT: P>0.05, CO: P>0.05, Table 3). The percent change in stem density is similar for both treatments: tillage in UT had an increase of 139% (sc, 2020: 19±7; 2022: 45±11) and in CO a decrease of 65% (sc, 2020: 93±35;2022: 31±39). In UT, tillage with *P. suaveolens* had a stem density increase of 147% (sc, 2020: 20±6; 2022: 48±18) and in CO a decrease of 67% (sc, 2020: 85±38; 2022: 28±24) (Table 2). In CO, tillage combined with *P. suaveolens* resulted in a slightly greater decrease in stem density in the first year (Figure 1) compared to tillage alone. The higher annual precipitation in the first year (Supplementary Table S1) may have contributed to *P. suaveolens* and tillage having a greater effect than the second year. In our study, application of *P. suaveolens* inoculum did not have a significant interaction with tillage but could be implemented in an IWM approach. Tillage has been used to manage Canada thistle by reducing stem density through the depletion of root reserves and reduction in shoot biomass (Thomsen 2011; Weigel 2024). Proper timing of tillage can be crucial, as early tillage can allow Canada thistle to recover and rebuild root reserves for overwintering (Donald 2000; Thomsen et al. 2015). Applying *P. suaveolens* inoculum two weeks after tillage may enhance pathogen invasion of the smaller root fragments, and increase systemic infection in the spring. (Alexopoulos et al. 1996; Berner et al. 2015a).

### Groundcover

#### Canada Thistle

The trends observed in Canada thistle cover align closely with those in stem density. When cover measurements are divided by stem density, an estimate of the biomass of individual shoots can be made which may indicate the health of the population. Treatments with herbicide had the lowest amount of Canada thistle cover, ranging from 0-25% in UT and 0-35% in CO. Of note, in 2022, plots treated with herbicide and *P. suaveolens* had zero Canada thistle cover while plots treated with herbicide alone still had a low density of Canada thistle stems. *Puccinia suaveolens* may have an additive effect in herbicide treated areas or help prevent regrowth, suggesting that these two treatments are compatible. Mowing also significantly decreased percent Canada thistle cover compared to control, although no significant difference occurred between mowing alone and mowing with *P. suaveolens*. The highest Canada thistle cover occurred in the control, which was more easily seen in UT than in CO (Figure 2). Thistle cover was higher in tilled plots as a result of the fragmentation and spread of Canada thistle roots creating many small populations (Donald 2000; Thomsen et al. 2015).

#### Vegetation

In UT, the herbicide treatment, which reduced broadleaf plants, including Canada thistle, allowed more opportunity for grasses to grow (≤ 80% cover in year 2). Grass cover in UT was significantly higher in herbicide treatments compared to control and tillage (UT: P<0.005), with greater effects observed in the second year. In a three year study of management tactics for a non-native forb, a positive effect on native grass cover was seen as a result of herbicide treatments, however a steady and significant increase in non-native grass cover was also seen (Skurski et al. 2013). Grasses were not identified to the species level and could include undesirable invasive species of concern (e.g., cheatgrass) for management of natural areas. In contrast, grass coverage in CO showed minimal change across treatments (P>0.05), though a significant interaction occurred between management and *P. suaveolens* (P=0.002). Further analysis showed that HMT and *P. suaveolens* treatments had significantly lower grass cover compared to control, *P. suaveolens*, or mowing plus *P. suaveolens* (Tables 4; Figure 2). Tillage to 20cm distributes seeds throughout the entire tillage area reducing accumulation of seeds near the surface, and therefore might result in reduced germination and grass cover as seen in our results (Feledyn-Szewczyk et al. 2020). Differing climatic conditions between the two sites could also affect grass growth and contribute to the difference between treatments. Therefore site characteristics need to be considered in conjunction with management strategy for potential revegetation or secondary invasion by non-native species.

Prior to initial treatments, forb cover was low (0-10%) and remained below 10% across the majority of plots with no significant difference between treatments (UT: P>0.05, CO: P>0.05) (Table 4, Figure 2). Use of broadleaf herbicides against invasive forbs can be expected to also suppress both native and other exotic forbs within the treatment areas (Skurski et al. 2013). As expected, only treatments with broadleaf selective herbicides showed a slightly greater decline in forb cover. In other studies, short-term changes in native forb cover remained insignificant after herbicide application, except for reductions in flowering and seed set for at least 4 years post-treatment (Crone et al. 2009). There may be long term implications in native forb recovery after herbicide is used to control non-native forbs like Canada thistle.

#### Non-Vegetation

Bare ground significantly increased as a result of HMT treatments with and without *P. suaveolens* in UT (P<0.001). However herbicide treatments, with and without *P. suaveolens* did not result in a significant difference compared to other treatments (P>0.05). Combined treatments had more of an impact on bare ground cover in UT than herbicide alone perhaps due to significantly more grass cover in the herbicide alone treated plots in year two (P<0.0001). Combined control tactics have been shown in both cropping and non-cropping systems to have better long-term control of Canada thistle than herbicide alone (Davis et al. 2018). There may have been an additional effect from changes in seedbank availability due to tillage (Feledyn-Szewczyk et al. 2020) or from mowing as mowing alone resulted in more bare ground compared to control (P=0.03). In CO, bare ground cover was significantly greater in both herbicide and HMT treated plots compared to all other treatments (P<0.001) (Table 4, Figure 2). There was no significant difference found between bare ground cover in HMT and herbicide alone treated plots. Bare ground is an important aspect of land management since it may necessitate reseeding to prevent soil erosion and the creation of niches for other noxious weeds. Revegetation should be included to promote native and desirable plants (Molvar et al. 2024; Rodriguez et al. 2024; Weidlich et al. 2020).

At the UT site, herbicide treatments resulted in significantly more litter compared to tillage (UT: P=0.002). In CO, HMT treatments had significantly lower litter compared to all treatments except for control (Table 4; Figure 2). Litter cover can be beneficial in reducing bare ground, retaining soil moisture and increasing nutrient cycling (Perera et al. 2024; Redman 1978)

### Practical Implications

Effective strategies for controlling Canada thistle vary with site conditions and management goals. Chemical treatments are often the most effective at reducing Canada thistle populations, but frequently increase areas of bare ground for invasion from other weeds and even re-invasion by Canada thistle. The use of herbicides is often restricted in natural landscapes or on organically certified farms, leaving stakeholders in search of alternative and complimentary control tactics. Applications of *P. suaveolens* inoculum in addition to other tactics show promising trends for greater control of Canada thistle than without the inoculum. Continued monitoring will help determine if additional treatment applications are needed, especially where *P. suaveolens* effects are enhanced on Canada thistle. Although *P. suaveolens* is slower acting in suppressing Canada thistle, one of the long-term benefits is that recovery of native plant communities is more likely. *Puccinia suaveolens* is unique among tactics for managing Canada thistle, as there are no direct effects on non-target plants. Applying *P. suaveolens* alone or with other tactics, provides a safe, sustainable, and effective means for enhancing management of Canada thistle in natural areas.

## Acknowledgments

The authors wish to thank, Sara Jo Dickens, PhD of Ecology Bridge LLC; Logan Jones of Park City Municipal Corporation, Levi Kokes Colorado Parks and Wildlife, Tamarack Ranch State Wildlife Area, Colorado State University Extension Office, and the Utah Weed Supervisors Association for site access, Park City site information signage, and other logistical and outreach support.

## Herbicide, mowing, tillage and rust application methods

**Table 1:** Overview of weed management tactics employed for treatment of Canada thistle at experimental sites in Colorado and Utah.

| Treatments | Equipment used | Application | Method | Optimal conditions and timing |
| --- | --- | --- | --- | --- |
| Inoculation with *P. suaveolens* | Portable one-gallon spray tank | 3.3g/m<sup>2</sup> inoculum | Wet all foliage with plain water, broadcast disperse inoculum, spray with water a second time. | Average daily temperature 65° F Fall.<br>Better with long dew periods such as evening. Added moisture layer. No wind |
| Herbicide (Chlorsulfuron and Aminopyralid tank mix) | Backpack 4-gallon manual piston-pump sprayer | Aminopyralid at 7 fl. oz/acre (0.11 acid equivalent), chlorsulfuron at 1 fl. oz/acre with non-ionic surfactant and marker dye tank mix | Broadcast spray plot directly after mowing if combined treatment. | Low wind, no rain, straight after mowing to help with translocation |
| Tillage | Honda 270cc Tiller Rear Tine 20" | Tilled to depth of 12 in. | Till 14 days post mowing or herbicide and directly preceding inoculations if combined. | Best when with other methods. Fall best timing. Earlier in the season requires more applications |
| Mowing | STIHL FS 91 16.5 in. Gas Trimmer | Mowed to height of 6 in. | Mow 2 weeks preceding inoculations. | Before seeding ideal |

## Percentage change of Canada thistle stem density Colorado and Utah

**Table 2:** Annual average change (percent) of Canada thistle and average stem count with +/- SD in Colorado and Utah from 2020-2022 following treatment with individual and combined weed management tactics. Based on a Tukey’s fence method for determining outliers, plots 74 and 75 from block 8 in CO were excluded from analysis.

| Treatment | Rust | Colorado |  |  |  |  |  | Utah |  |  |  |  |  |
| --- | --- | --- | --- | --- | --- | --- | --- | --- | --- | --- | --- | --- | --- |
|  |  | Fall<br>2020-20 | Stem<br>count<br>± | Fall<br>2021-20 | Stem<br>count<br>± | Fall<br>2020-20 | Stem<br>count<br>± | Fall<br>2020-20 | Stem<br>count<br>± | Fall<br>2021-20 | Stem<br>count<br>± | Fall<br>2020-20 | Stem<br>count<br>± |
|  |  | 21 %<br>change | 2020 | 22 %<br>change | 2021 | 22 %<br>change | 2022 | 21 %<br>change | 2020 | 22 %<br>change | 2021 | 22 %<br>change | 2022 |
| Control | NO | -3% | 76 ± 31 | -15% | 74 ± 28 | -18% | 63 ± 55 | 852% | 22 ± 16 | 7% | 40 ± 9 | 98% | 43 ± 19 |
| Control | YES | -16% | 93 ± 27 | -31% | 78 ± 36 | -42% | 54 ± 35 | 81% | 26 ± 8 | -19% | 47 ± 8 | 46% | 38 ± 13 |
| Tillage | NO | -26% | 93 ± 35 | -53% | 69 ± 27 | -65% | 32 ± 39 | 128% | 19 ± 7 | 5% | 43 ± 14 | 139% | 45 ± 11 |
| Rust, Tillage | YES | -48% | 85 ± 38 | -36% | 44 ± 39 | -67% | 28 ± 24 | 118% | 20 ± 6 | 13% | 43 ± 16 | 147% | 48 ± 18 |
| Mowing | NO | -39% | 83 ± 37 | -15% | 50 ± 30 | -48% | 43 ± 35 | 21% | 22 ± 9 | -18% | 26 ± 9 | -1% | 21 ± 11 |
| Rust, Mowing | YES | -25% | 62 ± 41 | -53% | 47 ± 36 | -65% | 22 ± 25 | 18% | 25 ± 17 | -26% | 30 ± 16 | -13% | 22 ± 8 |
| Herbicide | NO | -93% | 78 ± 21 | -77% | 6 ± 6 | -98% | 1 ± 2 | -97% | 29 ± 13 | -100% | 1 ± 2 | -100% | 0 ± 0 |
| Rust, Herbicide | YES | -90% | 84 ± 40 | -100% | 8 ± 9 | -100% | 0 ± 0 | -98% | 22 ± 14 | -100% | 1 ± 1 | -100% | 0 ± 0 |
| Mowing, Herbicide, Tillage | NO | -97% | 67 ± 44 | -93% | 2 ± 4 | -100% | 0 ± 0 | -69% | 25 ± 10 | -84% | 8 ± 5 | -95% | 1 ± 3 |
| Rust, Mowing, Herbicide, Tillage | YES | -86% | 84 ± 47 | -94% | 12 ± 15 | -99% | 1 ± 1 | -84% | 27 ± 7 | -10% | 4 ± 5 | -100% | 0 ± 0 |

## Canada thistle stem count ANOVA table Colorado and Utah

**Table 3:** Statistical results of the impact of rust inoculum application,, management practice, and their combination across seasons on Canada thistle stem count in Colorado and Utah.

|  | Utah |  |  | Colorado |  |  |
| --- | --- | --- | --- | --- | --- | --- |
|  | Chisq | Df | Pr(>Chisq | Chisq | Df | Pr(>Chisq |
| Rust | 0.0153 | 1 | 0.9017 | 0.0035 | 1 | 0.9017 |
| Management | 60.6082 | 4 | <0.0001 | 67.4382 | 4 | 2.161e-12 |
| Season | 0.7936 | 2 | 0.6725 | 138.1639 | 2 | 0.6725 |
| Management: Rust | 3.1770 | 4 | 0.5286 | 6.5493 | 4 | 0.5286 |
| Rust: Season | 1.1180 | 2 | 0.5718 | 2.6086 | 2 | 0.5718 |
| Management: Season | 170.0956 | 8 | <0.0001 | 117.6077 | 8 | < 2.2e-16 |
| Management: Rust: Season | 2.2153 | 8 | 0.9737 | 15.1340 | 8 | 0.9737 |

## Ground coverage ANOVA table for Colorado and Utah sites

**Table 4:**
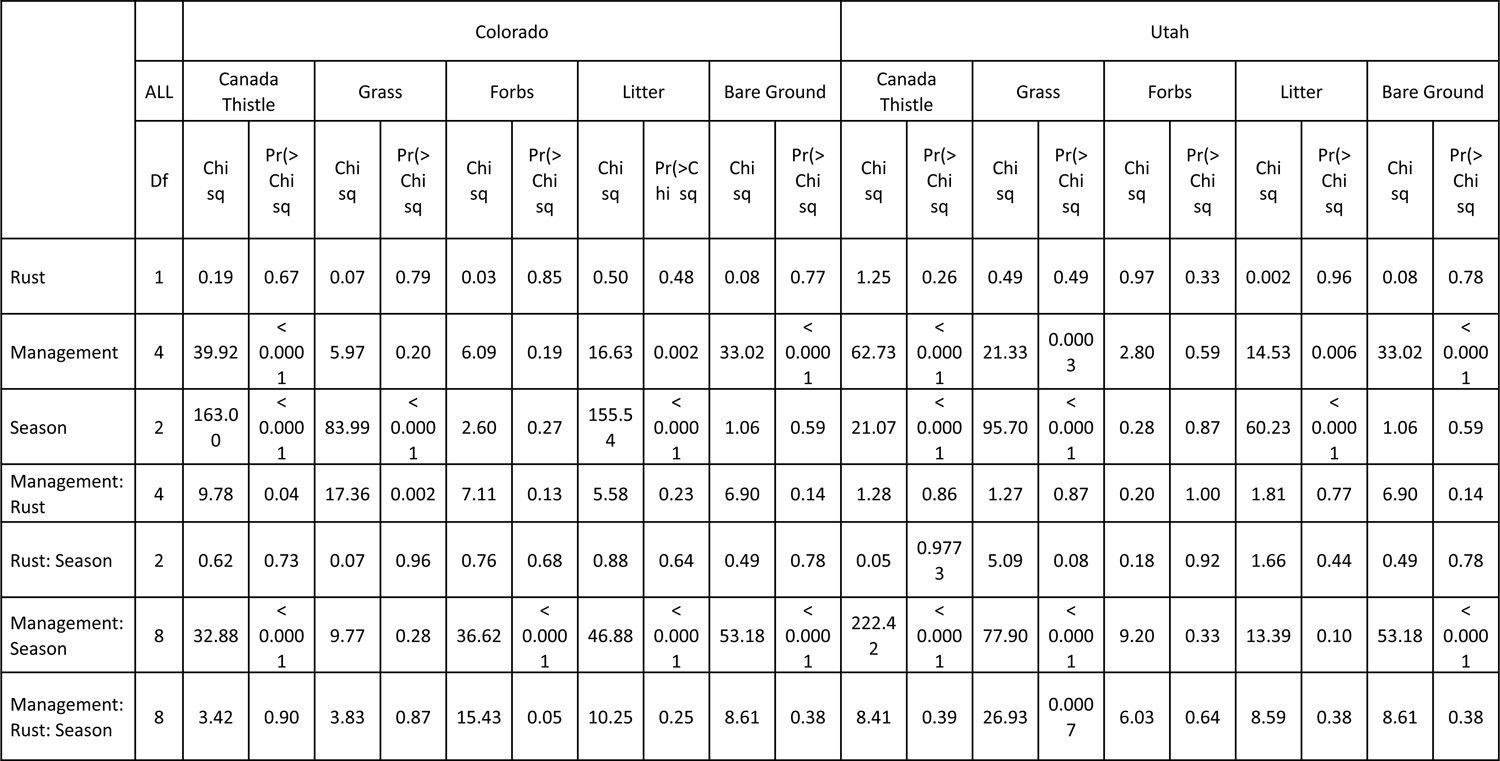
ANOVA table of the five ground cover types measured in UT and CO sites as a function of rust inoculum application, management strategy, season and the combined effects of these three parameters.

|  |  | Colorado |  |  |  |  |  |  |  |  |  | Utah |  |  |  |  |  |  |  |  |  |
| --- | --- | --- | --- | --- | --- | --- | --- | --- | --- | --- | --- | --- | --- | --- | --- | --- | --- | --- | --- | --- | --- |
|  | ALL | Canada Thistle |  | Grass |  | Forbs |  | Litter |  | Bare Ground |  | Canada Thistle |  | Grass |  | Forbs |  | Litter |  | Bare Ground |  |
|  | Df | Chi sq | Pr(> Chi sq | Chi sq | Pr(> Chi sq | Chi sq | Pr(> Chi sq | Chi sq | Pr(> Chi sq | Chi sq | Pr(> Chi sq | Chi sq | Pr(> Chi sq | Chi sq | Pr(> Chi sq | Chi sq | Pr(> Chi sq | Chi sq | Pr(> Chi sq | Chi sq | Pr(> Chi sq |
| Rust | 1 | 0.19 | 0.67 | 0.07 | 0.79 | 0.03 | 0.85 | 0.50 | 0.48 | 0.08 | 0.77 | 1.25 | 0.26 | 0.49 | 0.49 | 0.97 | 0.33 | 0.002 | 0.96 | 0.08 | 0.78 |
| Management | 4 | 39.92 | < 0.0001 | 5.97 | 0.20 | 6.09 | 0.19 | 16.63 | 0.002 | 33.02 | < 0.0001 | 62.73 | < 0.0001 | 21.33 | 0.0003 | 2.80 | 0.59 | 14.53 | 0.006 | 33.02 | < 0.0001 |
| Season | 2 | 163.00 | < 0.0001 | 83.99 | < 0.0001 | 2.60 | 0.27 | 155.54 | < 0.0001 | 1.06 | 0.59 | 21.07 | < 0.0001 | 95.70 | < 0.0001 | 0.28 | 0.87 | 60.23 | < 0.0001 | 1.06 | 0.59 |
| Management: Rust | 4 | 9.78 | 0.04 | 17.36 | 0.002 | 7.11 | 0.13 | 5.58 | 0.23 | 6.90 | 0.14 | 1.28 | 0.86 | 1.27 | 0.87 | 0.20 | 1.00 | 1.81 | 0.77 | 6.90 | 0.14 |
| Rust: Season | 2 | 0.62 | 0.73 | 0.07 | 0.96 | 0.76 | 0.68 | 0.88 | 0.64 | 0.49 | 0.78 | 0.05 | 0.9773 | 5.09 | 0.08 | 0.18 | 0.92 | 1.66 | 0.44 | 0.49 | 0.78 |
| Management: Season | 8 | 32.88 | < 0.0001 | 9.77 | 0.28 | 36.62 | < 0.0001 | 46.88 | < 0.0001 | 53.18 | < 0.0001 | 222.42 | < 0.0001 | 77.90 | < 0.0001 | 9.20 | 0.33 | 13.39 | 0.10 | 53.18 | < 0.0001 |
| Management: Rust: Season | 8 | 3.42 | 0.90 | 3.83 | 0.87 | 15.43 | 0.05 | 10.25 | 0.25 | 8.61 | 0.38 | 8.41 | 0.39 | 26.93 | 0.0007 | 6.03 | 0.64 | 8.59 | 0.38 | 8.61 | 0.38 |

## Canada thistle Stem count in Colorado and Utah

**Figure 1:**
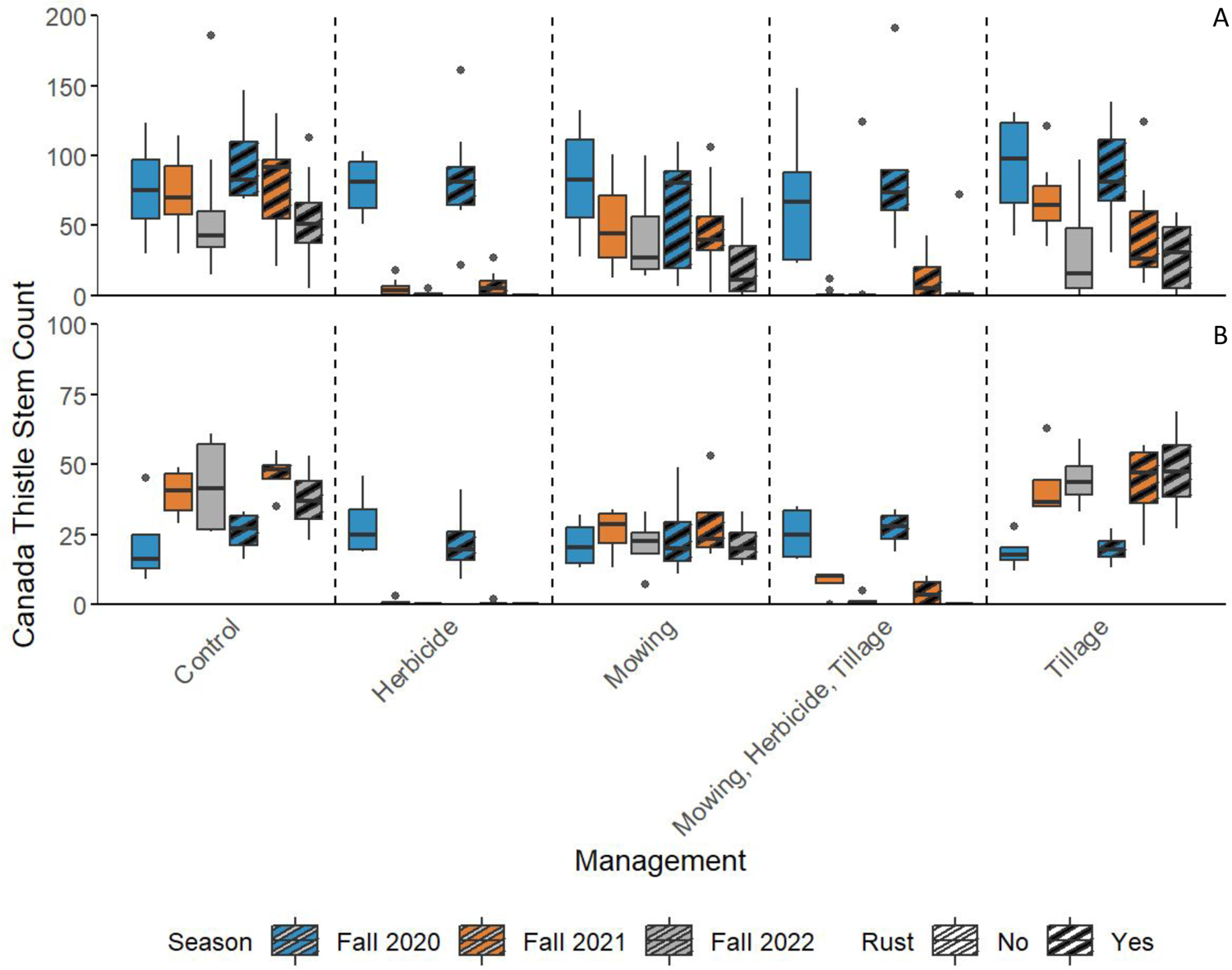
Canada thistle stem count in Fall 2020-2022 in A) Colorado and B) Utah following treatment with individual and combined weed management approaches.

## Ground percentage for weed management approaches at Colorado and Utah

**Figure 2:**
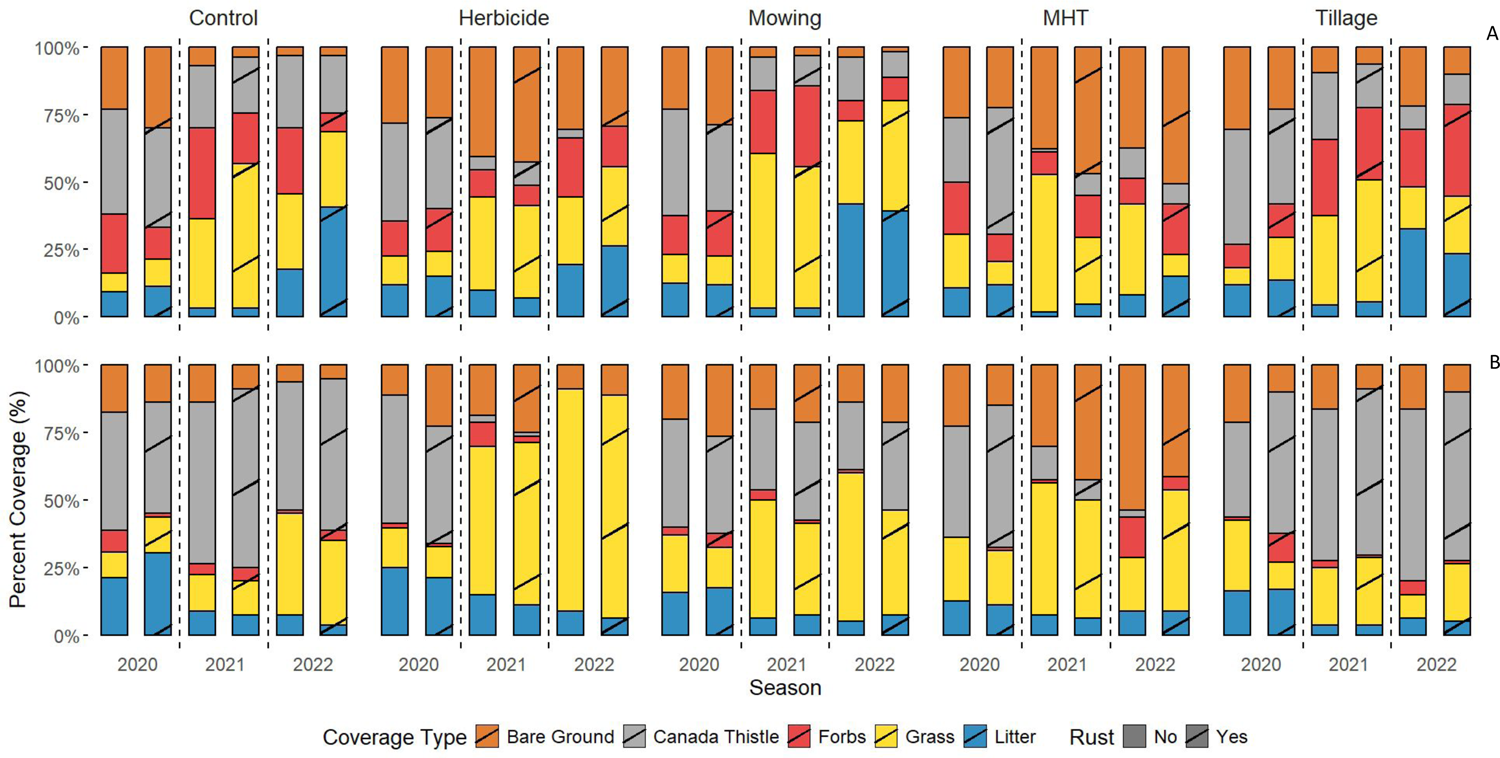
Average percent of the 5 measured ground cover types. A) Colorado and B) Utah experimental site, 2020-2022 following treatment with individual and combined weed management approaches.

## Climate table for Colorado and Utah 2020-2022

**Supplemental Table 1:** Precipitation, maximum and minimum temperatures at the two experiment sites (UT and CO) throughout the experimental years 2020-2022.

|  |  |  | Annual | Jan | Feb | Mar | Apr | May | Jun | Jul | Aug | Sep | Oct | Nov | Dec |
| --- | --- | --- | --- | --- | --- | --- | --- | --- | --- | --- | --- | --- | --- | --- | --- |
| CO | Precipitation (mm) | 2020 | 280.4 | 4.4 | 5 | 25.7 | 16.3 | 56.7 | 46.6 | 54.1 | 17.1 | 25.3 | 9.4 | 8.3 | 11.5 |
|  |  | 2021 | 488.1 | 1.7 | 11.2 | 65.7 | 50.2 | 108.2 | 56.1 | 67.5 | 93.4 | 12 | 14.6 | 6.6 | 0.8 |
|  |  | 2022 | 364.8 | 14.5 | 6.7 | 27.1 | 9.8 | 50.4 | 18.7 | 74.3 | 31.6 | 47.8 | 24 | 14.2 | 45.6 |
|  | Max Temperature (°C) | 2020 | 20.4 | 9.3 | 8.3 | 14.1 | 18.2 | 23 | 32.9 | 33.8 | 34.8 | 27.6 | 17.3 | 17 | 9 |
|  |  | 2021 | 19.9 | 5.8 | 1.7 | 14.1 | 16.6 | 22 | 31.4 | 34.2 | 32.7 | 30.5 | 21.7 | 16.5 | 11.2 |
|  |  | 2022 | 19.1 | 4.9 | 5.4 | 12 | 17.6 | 23.2 | 32.3 | 34.8 | 34.5 | 30.3 | 21.4 | 10.4 | 3.1 |
|  | Min Temperature (°C) | 2020 | 1.1 | -8.3 | -9.9 | -3 | -2.8 | 6.2 | 12.3 | 15.2 | 14.1 | 6.4 | -1.6 | -10 | -5.4 |
|  |  | 2021 | 1.5 | -9 | -13.5 | -4.3 | -1.5 | 6.3 | 12.6 | 15.1 | 13.3 | 9.2 | 1.7 | -3.8 | -8.5 |
|  |  | 2022 | 0.7 | -10.7 | -11.2 | -6.1 | -2.4 | 5 | 12.2 | 16.7 | 14.7 | 9.7 | 1.1 | -8.2 | -12.5 |
| UT | Precipitation (mm) | 2020 | 294.5 | 54.3 | 36.3 | 55.2 | 10.6 | 15.5 | 55.1 | 2.9 | 0.4 | 4.3 | 11.4 | 31.8 | 16.6 |
|  |  | 2021 | 612.1 | 34.5 | 59.3 | 44.8 | 42 | 33.2 | 10.4 | 31.6 | 109.9 | 11.3 | 111.6 | 20.4 | 103.1 |
|  |  | 2022 | 461.5 | 27.4 | 10.3 | 44 | 33.5 | 37.5 | 23.3 | 15.2 | 52.7 | 25.1 | 31.3 | 78.1 | 82 |
|  | Max Temperature (°C) | 2020 | 13.6 | 1.2 | 1.1 | 6.2 | 10.7 | 17.5 | 22.6 | 27.6 | 29.2 | 22.2 | 16.7 | 7 | 1.6 |
|  |  | 2021 | 13.4 | 1.2 | 0.4 | 5.2 | 10 | 17.2 | 26.6 | 28.7 | 24.7 | 22.1 | 12.9 | 10 | 1.8 |
|  |  | 2022 | 12.5 | 1.5 | 0.5 | 6.5 | 9.6 | 14 | 22.6 | 28.3 | 26.1 | 24.2 | 15.1 | 1.7 | -0.4 |
|  | Min Temperature (°C) | 2020 | 0.2 | -8 | -11.8 | -4.8 | -1.1 | 4.8 | 5.4 | 12.1 | 13.4 | 6.5 | -0.4 | -9.4 | 4.2 |
|  |  | 2021 | 0.4 | -10.7 | -8 | -5.2 | -2 | 3.7 | 9 | 14.7 | 9.7 | 5.4 | 0.6 | -3.9 | -8.1 |
|  |  | 2022 | -1.3 | -10.7 | -11.6 | -5.3 | -2.1 | -1.2 | 6.3 | 12.3 | 11 | 5.6 | -0.3 | -9.4 | -10 |

## References

Alexopoulos CJ, Mims CW, Blackwell M. (1996) Phylum Basidiomycota Order Uredinales The Rust. Pages 598-638 in Introductory Mycology. University of California: Wiley

Bean DW, Gladem K, Rosen K, Blake A, Clark RE, Henderson C, Kaltenbach J, Price J, Smallwood EL, Berner DK, Young SL, Schaeffer RN (2024) Scaling use of the rust fungus Puccinia punctiformis for biological control of Canada thistle (Cirsium arvense L. Scop.): First report on a U.S. statewide effort. Biological Control 192 105481, 10.1016/j.biocontrol.2024.105481

Beck KG, Sebastian JR (2000) Combining Mowing and Fall-Applied Herbicides to Control Canada Thistle (*Cirsium arvense*). Weed Technology 14(2):351–356, 10.1614/0890-037x(2000)014

Berner DK, Paxson LK, Bruckart WL, Luster DG, McMahon M, & Michael JL. (2002). First Report of Silybum marianum as a Host of Puccinia punctiformis. Plant disease, 86(11), 1271. 10.1094/PDIS.2002.86.11.1271B

Berner D, Smallwood E, Cavin C, Lagopodi A, Kashefi J, Kolomiets T, Pankratova L, Mukhina Z, Cripps M, Bourdôt G (2013) Successful establishment of epiphytotics of *Puccinia punctiformis* for biological control of *Cirsium arvense*. Biological Control 67(3):350–360, 10.1016/j.biocontrol.2013.09.010

Berner D, Smallwood E, Cavin C, McMahon M, Thomas K, Luster D, Lagopodi A, Kashefi J, Mukhina Z, Kolomiets T, Pankratova L (2015a) Asymptomatic systemic disease of Canada thistle (*Cirsium arvense*) caused by *Puccinia punctiformis* and changes in shoot density following inoculation. Biological Control 86:28–35, 10.1016/j.biocontrol.2015.02.006

Berner D, Smallwood E, Vanrenterghem M, Cavin C, Michael J, Shelley B, Kolomiets T, Pankratova L, Bruckart W, Mukhina Z (2015b). Some dynamics of spread and infection by aeciospores of *Puccinia punctiformis*, a biological control pathogen of Cirsium arvense. Biological Control 88:18–25, 10.1016/j.biocontrol.2015.05.001

Bodo Slotta T, Foley ME, Chao S, Hufbauer RA, Horvath DP (2010) Assessing Genetic Diversity of Canada Thistle (*Cirsium arvense*) in North America with Microsatellites. Weed Science 58(4):387–394, 10.1614/ws-d-09-00070.1

Bourdôt GW, Hurrell GA, Skipp RA, Monk J, Saville DJ (2011) Mowing during rainfall enhances the control of *Cirsium arvense*. Biocontrol Science and Technology 21(10):1213–1223, 10.1080/09583157.2011.

Bradshaw MJ, Carey J, Liu M, Bartholomew HP, Wayne M., Jurick WM II, Sarah Hambleton S, Hendricks D, Martin Schnittler M, Scholler M (2023) Genetic time traveling: sequencing old herbarium specimens, including the oldest herbarium specimen sequenced from kingdom Fungi, reveals the population structure of an agriculturally significant rust. New Phytologist 237: 1463–1473 doi: 10.1111/nph.18622

Brooks ME, Kristensen K, Benthem KJ, van Magnusson A, Berg CW, Nielsen A, Skaug HJ, Maechler M, Bolker BM (2017) {glmmTMB} Balances Speed and Flexibility Among Packages for Zero-inflated Generalized Linear Mixed Modeling. The R Journal 9(2):378– 400, 10.32614/RJ-2017-066

Buller AHR (1950) Puccinia suaveolens and its sexual process. Researches on fungi, Vol. VII: The sexual process in the Uredinales. University of Toronto Press, Toronto, pp. 344–388.

Carpinelli MF (2004) Autumn Mowing may Cut Herbicide Need. Agricultural Research 52(5):23, https://www.proquest.com/scholarly-journals/autumn-mowing-may-cut-herbicide-need/docview/208058950/se-2

Chichinsky D, Larson C, Eberly J, Menalled FD and Seipel T (2023) Impact of *Puccinia punctiformis* on *Cirsium arvense* performance in a simulated crop sequence. Front. Agron. 5: 1201600.doi: 10.3389/fagro.2023.1201600

Clark AL, Jahn CE, Norton AP (2020) Initiating plant herbivory response increases impact of fungal pathogens on a clonal thistle. Biological Control 143:104207, 10.1016/j.biocontrol.2020.104207

Connick WJ, French RC (1991) Volatiles emitted during the sexual stage of the Canada thistle rust fungus and by thistle flowers. J. Agric. Food Chem 39(1):185–188, 10.1021/jf00001a037

Cripps MG, Bourdôt GW, Saville DJ, Hinz HL, Fowler SV, Edwards GR (2011a) Influence of insects and fungal pathogens on individual and population parameters of *Cirsium arvense* in its native and introduced ranges. Biological Invasions 13(12):2739–2754, 10.1007/s10530-011-9944-7

Cripps MG, Gassmann A, Fowler SV, Bourdôt GW, McClay AS, Edwards GR (2011b) Classical biological control of *Cirsium arvense*: Lessons from the past. Biological Control 57(3):165– 174, 10.1016/j.biocontrol.2011.03.011

Cripps M, Bourdôt G, Saville DJ, Berner DK (2014) Success with the rust pathogen, Puccinia punctiformis, for biological control of Cirsium arvense. Pages 83-88 in XIV International Symposium on Biological Control of Weeds

Crone EE, Marler M, Pearson DE (2009) Non-target effects of broadleaf herbicide on a native perennial forb: a demographic framework for assessing and minimizing impacts. J. Appl. Ecol. 46 (3), 673–682.

Davis S, Mangold J, Menalled F, Orloff N, Miller Z, Lehnhoff E. A Meta-analysis of Canada Thistle (*Cirisium arvense*) Management. Weed Science 2018;66(4):548–57. doi:10.1017/wsc.2018.6

Demers A, Berner D, Backman P (2006) Enhancing incidence of *Puccinia punctiformis*, through mowing, to improve management of Canada thistle (*Cirsium arvense*). Biological Control 39(3):481–488, 10.1016/j.biocontrol.2006.06.014

Donald WW (2000) A degree-day model of *Cirsium arvense* shoot emergence from adventitious root buds in spring. Weed Science 48(3):333–341, 10.1614/0043-1745(2000)048

Enloe SF, Lym RG, Wilson R, Westra P, Nissen S, Beck G, Moechnig M, Peterson V, Masters RA, Halstvedt, M (2007). Canada Thistle (*Cirsium arvense*) Control with Aminopyralid in Range, Pasture, and Noncrop Areas. Weed Technology 21(4):890–894, 10.1614/wt-07-004.1

Feledyn-Szewczyk B, Smagacz J, Kwiatkowski CA, Harasim E, Woźniak A (2020) Weed Flora and Soil Seed Bank Composition as Affected by Tillage System in Three-Year Crop Rotation. Agriculture 10(5), 186, 10.3390/agriculture10050186

Fox J, Weisberg S eds (2019) An {R} Companion to Applied Regression. 3rd ed. Sage

French R, Lightfield A (1990) Induction of systemic aecial infection in Canada thistle (*Cirsium arvense*) by teliospores of *Puccinia punctiformis*. Phytopathology 80(9):872–877

Graglia E, Melander B, Jensen RK (2006) Mechanical and cultural strategies to control *Cirsium arvense* in organic arable cropping systems. Weed Research 46(4):304–312, 10.1111/j.1365-3180.2006.00514.x

Guske S, Schulz B, Boyle C (2004) Biocontrol options for *Cirsium arvense* with indigenous fungal pathogens. Weed Research 44(2):107–116, 10.1111/j.1365-3180.2003.00378.x

Harker KN, O’Donovan JT. Recent Weed Control, Weed Management, and Integrated Weed Management. Weed Technology. 2013;27(1):1–11. doi:10.1614/WT-D-12-00109.1

Hartig F, Lohse L (2024) Residual Diagnostics for Hierarchical (Multi-Level / Mixed) Regression Models [R package DHARMa version 0.4.7]. http://florianhartig.github.io/DHARMa/

Hettwer U, Gerowitt B (2004) An investigation of genetic variation in *Cirsium arvense* field patches. Weed Research 44(4): 289–297, 10.1111/j.1365-3180.2004.00402.x

Jacobs J (2006) Ecology and Management of Canada thistle [Cirsium arvense (l.) Scop.]. Invasive species Technical note. Bozeman, MT: U.S.. Department of Agriculture, Natural Resources Conservation Service

Kluth S, Kruess A, Tscharntke T (2003) Influence of mechanical cutting and pathogen application on the performance and nutrient storage of *Cirsium arvense*. Journal of Applied Ecology 40(2):334–343, 10.1046/j.1365-2664.2003.00807.x

Lalonde RG, Roitberg BD (1994) Mating System, Life-History, and Reproduction in Canada Thistle (*Cirsium arvense*; Asteraceae). American Journal of Botany 81(1):21, 10.2307/2445558

Lenth RV (2024) Estimated Marginal Means, aka Least-Squares Means [R package emmeans version 1.10.5]. https://cran.r-project.org/package=emmeans

Liang, C., Wang, P., Wang, Z., Zhao, N., Li, X., Li, J., Zhang, L., Meng, Q. and Yan, H., 2024. *Puccinia suaveolens* causing leaf rust on *Cirsium setosum* in China. Plant Disease, 108(3), p.815

Mendgen K, Hahn M (2002). Plant infection and the establishment of fungal biotrophy. Trends Plant Sci. 7, 352–356. doi: 10.1016/S1360-1385(02)02297-5

Menzies B (1953) Studies on the Systemic Fungus, *Puccinia suaveolens*, Ann. Bot. 17(4):551–568 10.1093/oxfordjournals.aob.a083369

Molvar EM, Rosentreter R, Mansfield D, Anderson GM (2024) Cheatgrass invasions: History, causes, consequences, and solutions. Hailey, ID: Western Watersheds Project, 128 pp.

Moore RJ (1975) THE BIOLOGY OF CANADIAN WEEDS: *Cirsium arvense* (L.) Scop. Canadian Journal of Plant Science 55(4):1033–1048, 10.4141/cjps75-163

Morin L, Brown JF, Auld BA (1992a) Effects of Environmental Factors on Teliospore Germination, Basidiospore Formation, and Infection of *Xanthium occidentale* by *Puccinia xanthii*. Phytopathology 82(12):1443, 10.1094/phyto-82-1443

Morin L, Brown JF, Auld BA (1992b) Teliospore germination, basidiospore formation and the infection process of *Puccinia xanthii* on *Xanthium occidentale*. Mycological Research 96(8):661–669, 10.1016/s0953-7562(09)80494-2

Morishita DW (1999) Canada Thistle. Pages 162-174 in Biology and Management of Noxious Rangeland Weeds. Corvallis, OR: Oregon State Press

Moyer JR, Schaalje GB, Bergen P (1991) Alfalfa (*Medicago sativa*) Seed Yield Loss Due to Canada Thistle (*Cirsium arvense*). Weed Technology 5(4):723–728. doi:10.1017/S0890037X00033753

Muir S, McClaran MP (1997) Rangeland inventory, monitoring, and evaluation. Arizona: Arizona Board of Regents, The Rangelands Partnership

Nadeau LB, Vanden Born WH (1989) The root system of Canada Thistle. Canadian Journal of Plant Science 69(4):1199–1206, 10.4141/cjps89-142

Nuzzo V (1997) ELEMENT STEWARDSHIP ABSTRACT for Cirsium arvense. Nature Conservancy.

Omernick, JM and Griffith, GE (2014) Ecoregions of the Conterminous United States: Evolution of a Hierarchical Spatial Framework. Environmental Management 54:1249–1266 doi: 10.1007/s00267-014-0364-1

Perera PCD, Gruss I, Szymura M (2024) Effect of litter decomposition on mowing and plant composition change during Solidago stand restoration. Ecological Questions 35(3):1–13, 10.12775/eq.2024.026

Rostrup E (1874) Eiendommeligt Generationsforhold hos Puccinia suaveolens (Pers.). Förhandlingar vid (ved) de Skandinaviska Naturforskare Mötet. 11:338–350

Petersen RH (1974) The rust fungus life cycle. The Botanical Review, 40(4):453–513, 10.1007/bf02860021

Peterson PG, Merrett MF, Fowler SV, Barrett DP, Paynter Q (2020) Comparing biocontrol and herbicide for managing an invasive non-native plant species: Efficacy, non-target effects and secondary invasion. Journal of Applied Ecology 57(10):1876–1884, 10.1111/1365-2664.13691

PRISM Climate Group (2023) Oregon State University https://prism.oregonstate.edu, data created 2020-2023, accessed 2020-2023.

O’Sullivan PA, Kossatz VC, Weiss GM, Dew DA (1982) An approach to estimating yield loss of Barley due to Canada Thistle. Canadian Journal of Plant Science 62(3):725–731, 10.4141/cjps82-105

R Core Team (2021). R: A language and environment for statistical computing. R Foundation for Statistical Computing. https://www.r-project.org/.

Redmann RE (1978) Plant and Soil Water Potentials following Fire in a Northern Mixed Grassland. Journal of Range Management 31(6):443, 10.2307/3897203

Rodriguez CS, McDonald CJ, Bean TM, Larios L (2024) Efficacy of invasive plant control depends on timing of herbicide application and invader soil seedbank density. Restoration Ecology 32(8), 10.1111/rec.14237

Sciegienka JK, Keren EN, Menalled FD (2011) Interactions between Two Biological Control Agents and an Herbicide for Canada Thistle (*Cirsium arvense*) Suppression. Invasive Plant Science and Management 4(1):151–158, 10.1614/ipsm-d-10-00061.1

Skurski TC, Maxwell BD, Rew LJ (2013) Ecological tradeoffs in non-native plant management. Biol. Conserv. 159, 292–302.

Thomas RF, Tworkoski TJ, French RC, Leather GR (1994) *Puccinia punctiformis* Affects Growth and Reproduction of Canada Thistle (*Cirsium arvense*). Weed Technology 8(3):488–493, 10.1017/s0890037X00039567

Thomsen MG, Mangerud K, Riley H, Brandsæter LO (2015) Method, timing and duration of bare fallow for the control of *Cirsium arvense* and other creeping perennials. Crop Protection 77:31–37, 10.1016/j.cropro.2015.05.020

Tiley GED (2010) Biological Flora of the British Isles: *Cirsium arvense* (L.) Scop. Journal of Ecology 98(4):938–983, 10.1111/j.1365-2745.2010.01678.x

Van Den Ende G, Frantzen J & Timmers T. Teleutospores as origin of systemic infection of *Cirsium arvense* by *Puccinia punctiformis*. Netherlands Journal of Plant Pathology 93, 233– 239 (1987). 10.1007/BF01998251

Weidlich EWA, Flórido FG, Sorrini TB, Brancalion PHS (2020) Controlling invasive plant species in ecological restoration: A global review. Journal of Applied Ecology 57(9):1806–1817, 10.1111/1365-2664.13656

Wickham H, Averick M, Bryan J, Chang W, McGowan L, François R, Grolemund G, Hayes A, Henry L, Hester J, Kuhn M, Pedersen T, Miller E, Bache S, Müller K, Ooms J, Robinson D, Seidel D, Spinu V, Takahashi K, Vaughan D, Wilke C, Woo K, Yutani H (2019) Welcome to the Tidyverse. The Journal of Open Source Software 4(43):1686, 10.21105/joss.01686

Wilson CL (1969) Use of plant pathogens in weed control. Annual Review of Phytopathogology. 7:411–434. 10.1146/annurev.py.07.090169.002211

